# Reconstructing cell interactions and state trajectories in pancreatic cancer stromal tumoroids

**DOI:** 10.1101/2022.02.14.480334

**Authors:** Ryo Okuda, Bruno Gjeta, Doris Popovic, Ashley Maynard, Qianhui Yu, Zhisong He, Malgorzata Santel, Makiko Seimiya, Soichiro Morinaga, Yohei Miyagi, Tomoyuki Yamaguchi, Yasuharu Ueno, Hideki Taniguichi, Barbara Treutlein, J. Gray Camp

**Affiliations:** Department of Regenerative Medicine, Yokohama City University Graduate School of Medicine, Kanagawa, Japan; Division of Regenerative Medicine, Center for Stem Cell Biology and Regenerative Medicine, the Institute of Medical Science, the University of Tokyo, Tokyo, Japan; Institute of Molecular and Clinical Ophthalmology Basel, Switzerland; University of Basel, Basel, Switzerland; Roche Institute for Translational Bioengineering (ITB), Roche Pharma Research and Early Development, Roche Innovation Center Basel, Switzerland; Department of Biosystems Science and Engineering, ETH Zürich, Basel, Switzerland; Department of Hepato-Biliary and Pancreatic Surgery, Kanagawa Cancer Center, Kanagawa, Japan; Molecular Pathology and Genetics Division, Kanagawa Cancer Center Research Institute, Kanagawa, Japan; Laboratory of Regenerative Medicine, School of Life Sciences, Tokyo University of Pharmacy and Life Sciences, Tokyo, Japan

**Author notes:** **Correspondence:** Hideki Taniguichi, J. Gray Camp, Barbara Treutlein. These authors contributed equally (Ryo Okuda, Bruno Gjeta).

## Abstract

Interlineage communication within a cancer microenvironment can augment cancer cell behaviour and impact response to therapy. Patient-derived cancer organoids provide an opportunity to explore cancer cell biology, however it is a major challenge to generate a complex cancer microenvironment in vitro. Here, we established a stromal tumoroid culture system modeling pancreatic ductal adenocarcinoma (PDAC) that reconstitutes multilineage interactions between cancer, endothelial, and fibroblast cells and recapitulates several aspects of primary tumors. Whole-mount immunohistochemistry on cleared tumoroids reveals organized vessel, desmoplastic fibroblast, and glandular cancer cell phenotypes that emerge over time. Time-course scRNA-seq measurements show that tumoroid formation activates fibroblasts, altering the extracellular matrix (ECM) composition and inducing cancer cell signal-response signatures and metabolic state change. Comparison between tumoroids with normal or cancer associated fibroblasts (CAFs) reveals different ECM compositions, as well as differential effects on cancer cell behaviors and metabolism. We identify Syndecan 1 (SDC1) and Peroxisome proliferator-activated receptor gamma (PPARG) as receptor and metabolic nodes involved in cancer cell response to CAF signals, and blocking SDC1 disrupts cancer cell growth within the tumoroid. Tumoroids from multiple PDAC patients revealed co-existence of subpopulations associated with classical and basal phenotypes, and CAF-induced migration behaviors emerged in certain patient tumoroids. Comparisons between patient tumoroids revealed a multigene migration signature that develops over time reflecting a stress response mechanism that correlates with worse clinical outcome. Altogether, stromal tumoroids can be used to explore dynamic and reciprocal interactions between cancer, CAF and endothelial cell states, and our data provides new inroads into the discovery of personalized pancreatic cancer therapies.

## Main text

Pancreatic ductal adenocarcinoma (PDAC) is one of the most aggressive and intractable forms of cancer [1]. PDAC tumors are characterized by stark intra-tumoral heterogeneity with a dense stroma component which can constitute over 70% of the tumor mass. Intratumoral heterogeneity in the tumor microenvironment (TME) originates in definable regional tissue states, and underlying sub-tumor microenvironments shape cancer phenotypes and can influence key clinical metrics of disease progression [2,3]. Cancer associated fibroblasts (CAFs) are central PDAC stroma components and coordinate diverse features of the TME including secreting cytokines that regulate cancer growth and shape evolutionary pressures that support malignancy [4].

Microenvironmental pressures within the primary tumor can lead individual cancer cells to acquire specific metabolic and other cell state signatures that support cancer cell adaptation to current conditions, and also provide the ability for future colonization into other organ niches [5–7]. These diverse pressures within the tumor are spatially and temporally dynamic, and it has been difficult to understand how CAF-cancer cell interactions generate a diversity of cell states. Cancer cystic organoids (CCOs) can be established from patients and provide extraordinary opportunities to study cancer cell biology [8,9]. Co-culturing cancer cells and CAFs in vitro is starting to provide new insights into how fibroblasts and cancer cells interact [10,11]. Single-cell sequencing enables the reconstruction of cell state continuums within complex developing tissues [12], and provides predictions for how cells interact based on analysis of receptor and ligand expression patterns between cell types [13]. Here, we set out to establish a stroma-rich tumoroid co-culture system to understand PDAC cancer-CAF interactions in controlled environments, and to explore developmental processes within tumoroids using single-cell transcriptome sequencing. By doing so, we learn about principles associated with intratumoral heterogeneity that might be leveraged for therapy development in PDAC and other cancers.

We generated cancer cyst organoids (CCOs) from PDAC patient primary biopsies, which we stably cultured according to previously published protocols [14] (Supplementary Table 1). Single-cell RNA sequencing (scRNA-seq) revealed that cell cycle state heterogeneity was a predominant source of variation in CCO cultures (Extended Data Fig. 1a-h). Comparison of CCO gene expression to matched bulk healthy and cancerous pancreatic tissue, revealed low correlation to healthy tissue, high correlation to pancreatic cancer tissue, and detection of PDAC-associated expression signatures within the CCOs (Extended Data Fig. 1i-l). To explore the developmental dynamics that underlie PDAC heterogeneity, we established a stroma-rich PDAC tumoroid co-culture system that combines cancer cells, endothelial cells (ECs) and CAFs grown in a three-dimensional matrix (Fig. 1a). We note that the CAFs and ECs were not derived from the same patient, and the same CAF and EC lines were used throughout our study. Over 48 hours, the cells form a spherical culture (Extended Data Fig. 2a), and over a 14 day period we observed substantial cellular organization that emerged within the tumoroid. Cancer cells form glandular structures, endothelial cells assemble into vessel networks, and CAFs produce extracellular matrix (Fig. 1b, Extended Data Fig. 2b-c). We used scRNA-seq to analyze heterogeneity in the tumoroids at day 7 and 14 of co-culture and compared the cell states with CCO, EC, and CAF mono-cultures (day 0) (Fig. 1c–d). The cells could be grouped into 12 clusters, representing CAF (clusters 1-5), EC (cluster 6,7), and cancer cell (clusters 8-12) states, and we provide marker genes for each cluster (Supplementary Table 2). Differential expression analysis between the 2D mono-culture and 3D tumoroid counterparts at day 7 and day 14 revealed diverse changes that emerged in the tumoroid over time (Extended Data Fig. 2d-g). There was a general hypoxia response for all cell types within the day 7 tumoroid, followed by angiogenic induction, extracellular component modulation, and metabolic adaptation signatures for the different cell types by day 14 (Supplementary Table 3). Strikingly, we found that tumoroid CAFs induced a consortium of extracellular matrix proteins (Collagens COL3A1, COL6A2, COL1A2, COL1A1; Fibronectin 1, FN1; Decorin, DCN) compared to the 2D CAFs, and tumorid CAF-enriched genes had gene ontology (GO) enrichments for transforming growth factor beta (TGF-β) signaling, inflammatory response, angiogenesis, and hypoxia. These data show that the multilineage tumoroid microenvironment induces strong morphological and molecular cell state changes across different cell types.

**Fig. 1.**
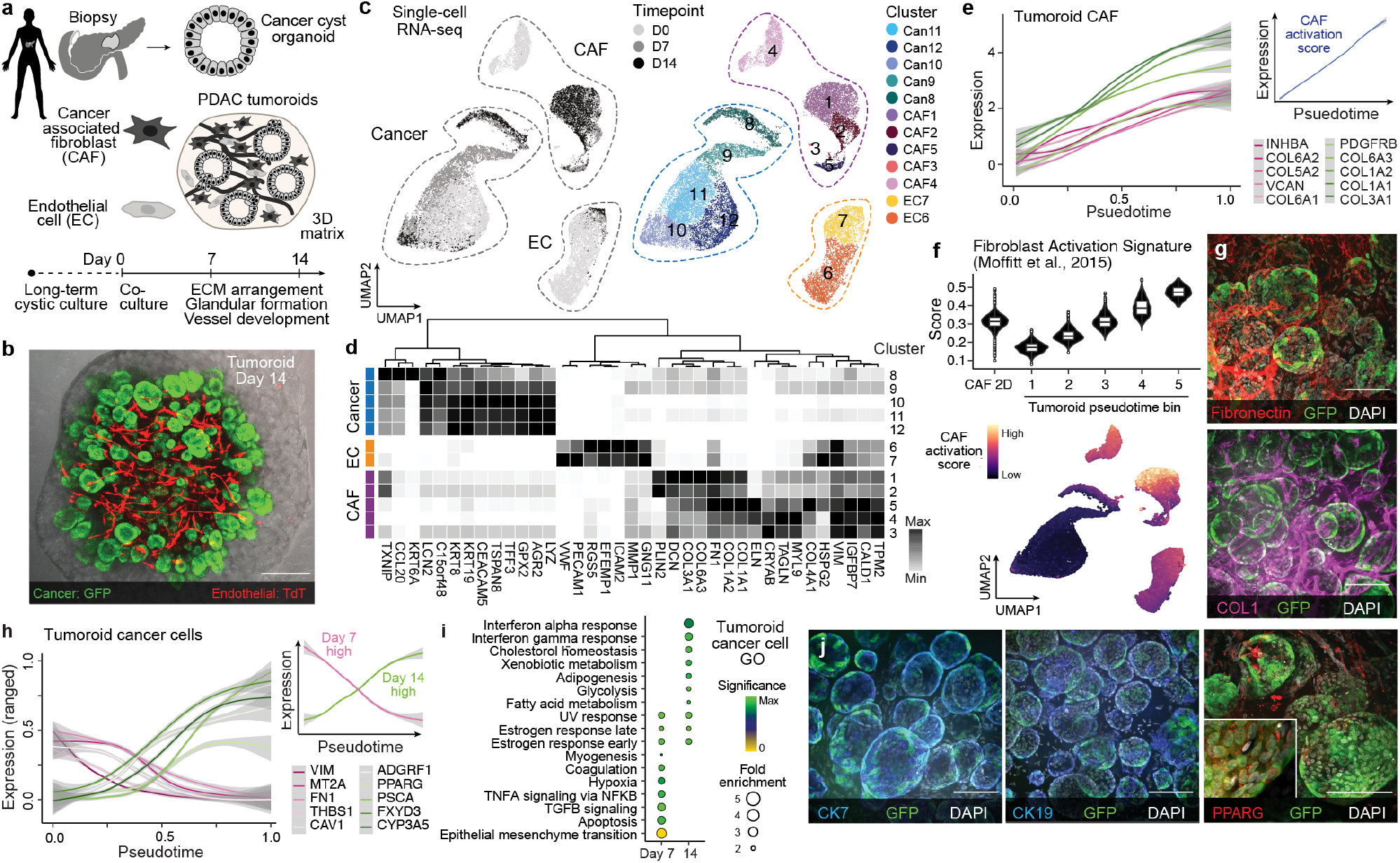
Reconstruction of cell state development within multilineage PDAC tumoroids using single-cell transcriptomics. a) Long-term cancer cyst organoid (CCO) cultures established from patients with pancreatic adenocarcinoma (PDAC) can be co-cultured in a 3D collagen/matrigel matrix with endothelial cells (EC) and cancer associated fibroblasts (CAF), which self-assemble a complex tumoroid microenvironment. Over 14 days, fibrous connective tissue forms, vessels sprout and organize, and cancer cells form 3D glandular structures within multilineage tumoroids. b) Day 14 tumoroid with cancer cells and ECs stably transformed with EGFP and TdT expression cassettes, respectively. Scale bar:250um. c) scRNA-seq was performed on the input cells in mono-culture (Day 0) and on tumoroids after 7 and 14 days in co-culture. UMAP cell embedding of scRNA-seq data is colored by time point (left) and by cluster (right). d) heatmap showing normalized expression of cluster marker genes. e) Expression of extracellular matrix protein encoding and other genes in CAFs ordered based on pseudotime reconstruction. Inset shows a CAF activation score based on primary PDAC cancer tissues15. f) Distributions of CAF activation score in mono-culture CAFs and tumoroid CAFs in 5 pseudotemporal bins (top) and UMAP with cells colored by Moffit et al. CAF activation score (bottom). g) Whole-mount immunohistochemistry on cleared tumoroids stained for extracellular matrix proteins (Fibronectin, Collagen 1). Scale bar:100um. Cancer cells stably express GFP, DAPI marks nuclei (white). h) Pseudotemporal expression profile of genes differentially expressed (DE) between tumoroid cancer cells at day 7 and 14. Inset shows cumulative expression profiles of the DE genes. i) Hallmark enrichment analysis on day 7 and 14 cancer cells. j) Whole-mount immunohistochemistry on cleared PCOs stained for cancer cell markers (CK17 and CK19, teal) and tumoroid-upregulated gene PPARG (red). Cancer cells stably express GFP, DAPI marks nuclei (white). Scale bar:100um.

To explore the effect of CAF signals on cell state dynamics within the tumoroid, we first established a pseudotemporal trajectory for each cell type and identified g enes t hat vary over the trajectory (Fig. 1e–f, Extended Data Fig. 2h-j). We observed that there were cells from both time points that covered the entire range of the trajectory, and for CAF we observed similar proportions of day 7 and day 14 CAFs along the CAF trajectory. In contrast, for cancer and endothelial cells there was a strong relationship between time point and position on the trajectory for both cell types. We observed that the CAF pseudotemporal ordering reflected activation status, such that induction of collagens and cytokines could be observed along pseudotime (Fig. 1e). Comparison to data from primary PDAC tissue revealed that the CAF temporal trajectory strongly resembles a transition from “normal” to “activated” stroma signatures [15] (Fig. 1f). This activated stroma signature has previously been shown to be prognostic, associating with worse clinical outcomes, and is characterized by the expression of genes that point to the role of CAF activation in tumor promotion. The CAF activation signature includes Secreted protein acidic and cysteine rich (SPARC), Wnt pathway family members (WNT5A and WNT5B), Matrix metalloproteinases (MMP2, MMP11 and MMP14), and Fibroblast Activation Protein (FAP)[15]. (Extended Data Fig. 2). Regulome analysis using SCENIC [16] suggests Early Growth Response 1 (EGR1) as a central transcriptional regulator that likely coordinates CAF activation (Extended Data Fig. 3a-c), that is upstream of several growth factor signaling pathways (BMP2, NOTCH3, LIF, VEGFC) and ECM regulators (COL5A3, COL12A1, LAMA4, HAS2) (Extended Data Fig. 3d-e). Whole-mount immunohistochemistry on cleared tumoroids stained for Collagen 1 and Fibronectin which revealed substantial ECM deposition surrounding cancer cells (Fig. 1g, Extended Data Video 1).

Along the EC trajectory, there was increased expression of Matrix Gla Protein (MGP), Angiopoietin-2 (ANGPT2), Endothelial cell-specific molecule 1 (ESM1) and other signatures of hypoxia response, angiogenesis, and TNF-α signalling that increase over pseudotime and have highest expression in day 14 tumoroids (Extended Data Fig. 2i-j). The cancer cell trajectory revealed initial induction of hypoxia-, apoptosis-, and epithelial-to-mesenchymal transition-related genes followed by adaption expression signatures associated with xenobiotic metabolism, cholesterol homeostasis, and interferon responses (Fig. 1h-j, Extended Data Video 2). Many of the genes that increase over pseudotime and peak at day 14 in tumoroid cancer cells, such as Peroxisome proliferator-activated receptor gamma (PPARG), Syndecan 1(SDC1), Mucin 1 (MUC1), Kruppel-like factor 3 (KFL3), have been previously associated with poor disease outcome [17–19]. Regulome analysis revealed transcription factors and their predicted targets that are differential along the cancer pseudotime, and these analyses highlighted a predominant role of PPARG and KLF2/3 in coordinating the cancer cell responses within the tumoroid (Extended Data Fig. 4a-c). Interestingly, the PPARG regulome linked fatty acid and cholesterol metabolism, with the IL2-STAT5, P53, and the Interferon signaling pathways (Extended Data Fig. 4d). In addition, PPARG correlated genes are also strongly associated with an immuno-suppressive environment, as well as lower survival in the PDAC tumor cohort from the cancer genome atlas (TCGA) (Extended Data Fig. 4e-h) [20]. Altogether, these data show that CAFs become activated by day 7 within the tumoroid, and suggest that CAF activation induces endothelial and cancer cell hypoxic and metabolic transition response states that are relevant for primary pancreatic cancer progression.

We next wanted to understand the specificity o f t he CAF-derived signals and their impact on cancer cell states. We generated tumoroids composed of CAF or normal fibroblasts (NF, derived from healthy pancreatic tissue), and analysed tumoroids using imaging and single-cell transcriptomics at day 7 and day 14 (Fig. 2a). We observed CAF tumoroids had larger glandular structures and more developed endothelial networks compared to NF tumoroids suggesting a difference in signalling cues between the two microenvironments (Extended Data Fig. 5a,b). Single-cell transcriptome analysis of CAF- or NF-cancer tumoroids revealed 4 cancer cell clusters (c3,4,6,8) and 5 fibroblast clusters (c0,1,2,5,7; Fig. 2b). We found that within the fibroblasts clusters, there were differential proportions of CAFs and NF (Fig. 2c). Clusters 1 and 2 were predominantly composed of CAF cells, conversely cluster 0 was enriched for NF cells (chisquared test p-value < 2.2e-16). Differentially expressed genes between NF and CAF show enrichments in both cases for epithelial-to-mesenchymal transition gene ontology, and CAF-enriched genes have increased enrichment for genes associated with angiogenesis, estrogen response, and the TP53 pathway (Fig. 2d,e; Extended Data Fig. 5c-e). We combined cancer cells from NF and CAF tumoroid conditions from both time points, and inferred cancer cell trajectories. We observed segregation of cells along the pseudotime trajectory that was time dependent, such that day 14 cancer cells grown with CAFs were predominantly enriched at later stages of trajectory (Fig. 2f). We searched for genes that increased along the trajectory, and were therefore enriched in day 14 CAF tumoroids, and observed upregulation of genes involved in metabolic homeostasis as well as many classical PDAC associated genes including TFF3, TSPAN8, AGR2, and multiple S100 (Fig. 2g, Extended Data Fig. 5f-h). Interestingly, we identify Lipocalin 2 (LCN2) as an early induced gene, and potentially could serve as an early prognostic biomarker of subsequent cancer metabolic state response to activated CAF signalling (Fig. 2h). We validated low expression of AKAP12 and CAF-specific induction of PDAC signature genes TFF3 and LCN2 using immunohistochemistry in tumoroids (Fig. 2i–k). Together, these data suggest that CAF-specific signals induce relevant cancer cell states, and that this interaction dynamic can be recapitulated and studied in developing tumoroid culture systems.

**Fig. 2.**
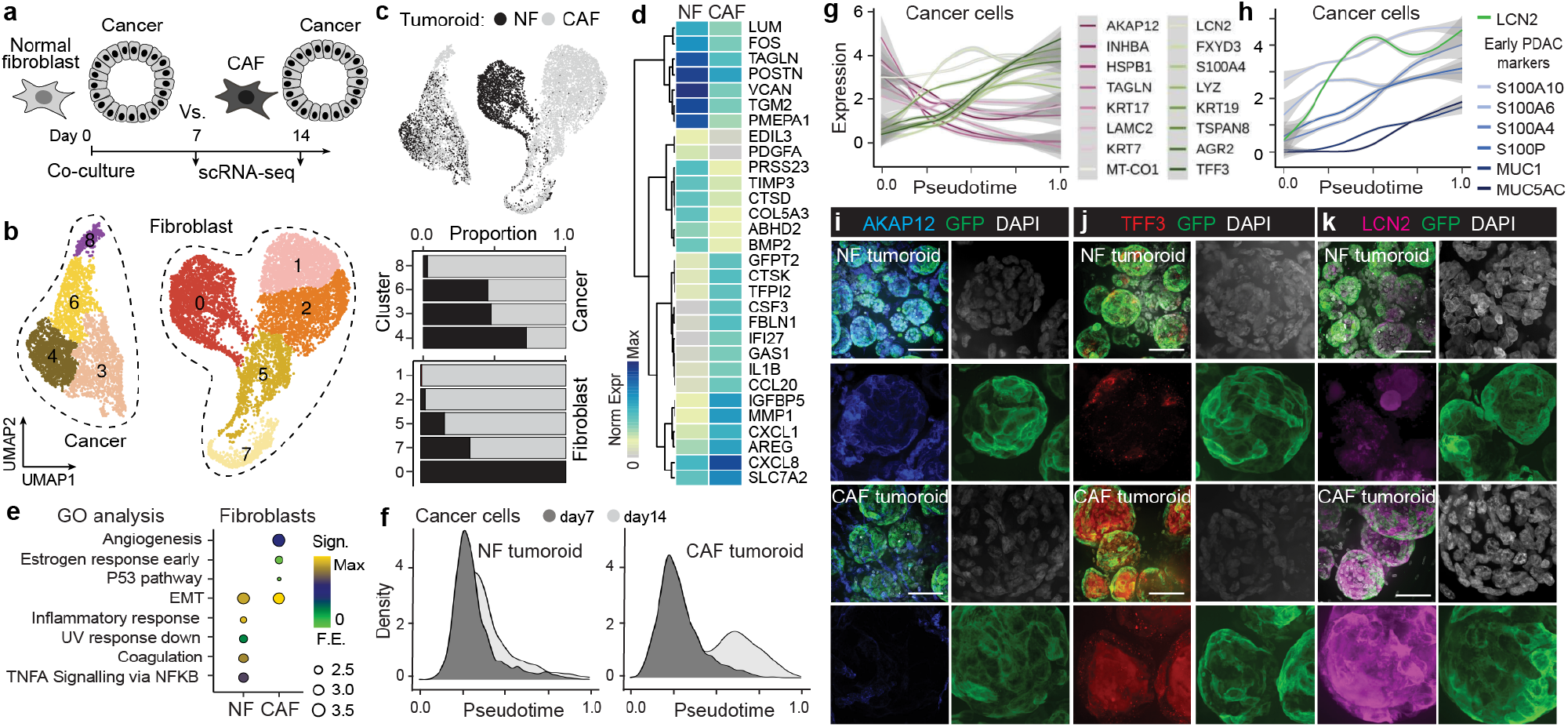
CAFs provide distinct signals from normal fibroblasts and promote cancer cell state change in tumoroids. a) Tumoroids containing normal or cancer associated fibroblasts were generated and analysed by scRNA-seq. b) UMAP embedding colored and numbered by cluster, with cancer and fibroblast cells encircled and noted. c) UMAP with cells colored by tumoroid type (top). Stacked barplot shows proportion of cancer or fibroblast cells per cluster and colored by tumoroid type. Clusters are significantly enriched for tumoroid type (chi-squared p-value < 2.2e-16). d) Heatmap shows expression of genes differentially expressed (DE) by NF and CAF in the tumoroids. e) Gene ontology enrimentments for NF and CAF DE genes. f) Density plots showing proportion of cancer cells along an inferred pseudotime in the NF and CAF tumoroids. Day 14 cancer cells have altered profiles only in CAF tumoroids. g-h) Expression profiles of genes over cancer cell pseudotime that are DE between day 7 and day 14. Day 14 DE genes are also DE between cancer cells in NF and CAF tumoroids. i-k) Immunofluorescence of AKAP12 (i), TFF3 (j), and LCN2 (k) protein expression in NF and CAF tumoroids. Cancer cells stably express GFP, DAPI marks nuclei (white). Scale bar:100um.

We used ligand-receptor pairing analyses to explore how CAF and cancer cells interact within the tumoroid (Extended Data Fig. 6a-d). We found that a substantial portion of CAF secreted signals are predicted to interact with the cancer expressed surface protein Syndecan 1 (SDC1) (Fig. 3a). SDC1 expression, as well as the predicted ligands, increase over time in the tumoroid (Fig. 3b) and it was previously revealed that SDC1 is recycled to the cell membrane by KRAS activity and is a critical mediator of macropinocytosis in pancreatic cancer [18].

**Fig. 3.**
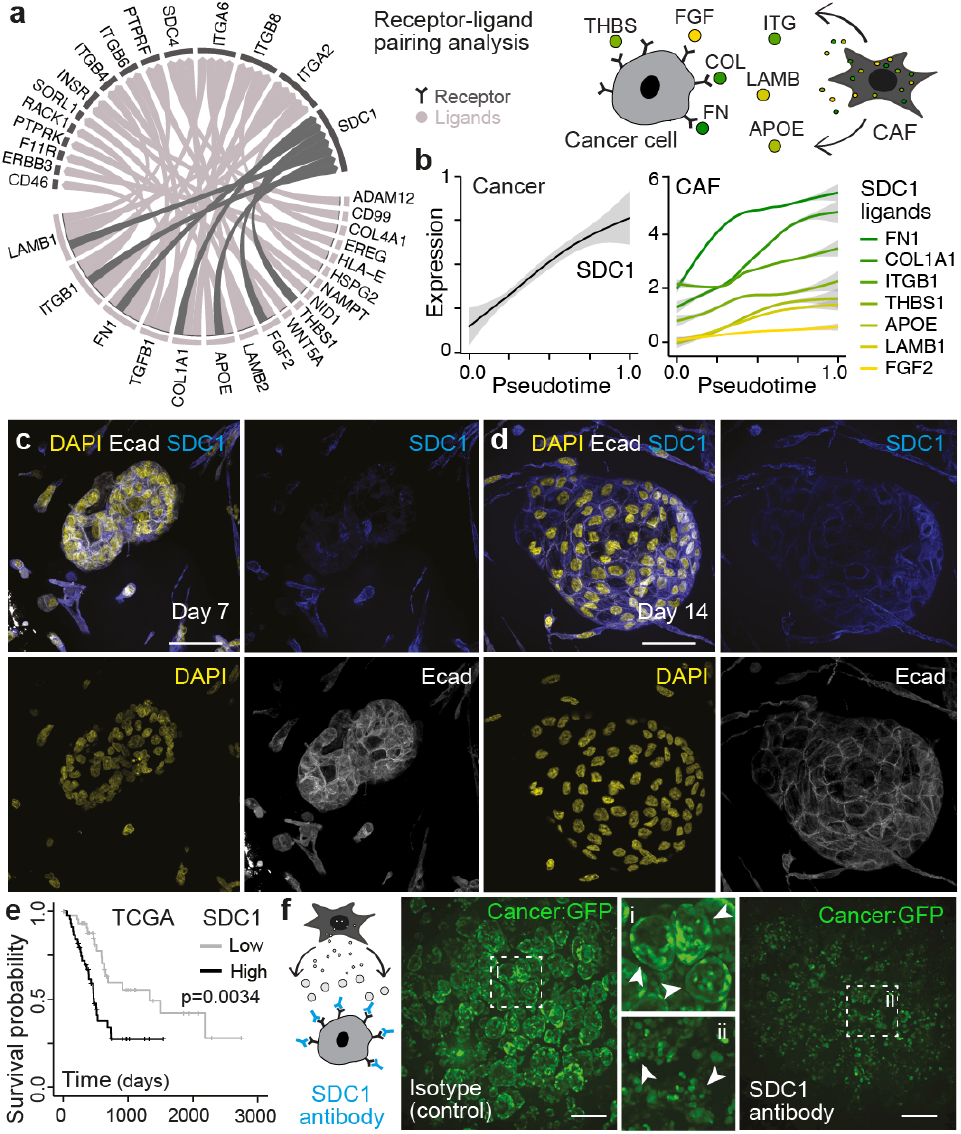
Interaction analysis identifies SDC1 as a dynamically expressed receptor in tumoroid cancer cells and SDC1-antibody block disrupts cancer cell glandular structure. a) Ribbon plot32 showing receptor-ligand pairing in CAF tumoroids. Syndecan 1 (SDC1)-ligand pairings are highlighted in dark grey. b) SDC1 (left) and predicted ligand (right) expression in cancer and CAF cells over pseudotime, respectively. c,d) Immunofluorescence showing SDC1 induction from day 7 (c) to day 14 (d) tumoroids. Scale bar:50um. e) Kaplan–Meier plot showing that high expression of SDC1 in PDAC cancers from the cancer genome atlas (TGCA) dataset is associated with lower survival. f) Schematic shows SDC1 antibody blocking experiment. GFP reporter expression in cancer cells on day 14 tumoroids incubated with isotype control (left, inset i) or SDC1 (right, inset ii) antibodies. Scale bar:100um.

SDC1 is known to be a key cell surface adhesion molecule engaged in interactions with numerous ligands (eg THBS1,FGF2,TNC,FN1) [21–26], thereby regulating major pathways responsible for cell interactions with the microenvironment, and contributing to cancer progression, proliferation, metastasis and overall poor prognosis [27]. We also searched for CAF receptors that might receive signals from cancer cells, and identified epidermal growth factor receptor (EGFR) as binding partner to cancer ligands that is associated with CAF activation in the tumoroid (Extended Data Fig. 6c-d). Primary PDAC tumors with high EGFR expression are associated with poor survival (Extended Data Fig. 6e), and EGFR inhibition shows promise as a co-target in mouse models and is FDA approved for PDAC treatment in humans [28,29]. More broadly, we show that regulators of diverse signalling pathways are dynamically modulated in the tumoroid (Extended Data Fig. 6g-f), providing a rich resource for future in vitro perturbation experiments to understand these complex interactions. Our data revealed SDC1 to be one of the highly enriched receptors, which prompted us to investigate how the blockage of SDC1 impacts cancer growth within the tumoroid. Immunofluorescence confirmed that SDC1 protein increases in expression in day 14 relative to day 7 CAF tumoroids, and is predominantly localized to the cancer cell membrane (Fig. 3c-d). We analyzed PDAC samples from the cancer genome atlas (TCGA) [20] and found that high detection of SDC1 is associated with lower overall survival (Fig. 3e). We cultured day 14 CAF tumoroids with an antibody blocking the activity of SDC1, and found substantial disruption of cancer cell growth (Fig. 3f). These data show that multilineage tumoroids can be used to manipulate and understand CAF-cancer interactions with therapeutic relevance.

To understand the potential diversity of molecular profiles and cell behaviors between different patients, we established tumoroids containing cancer cells from four additional patients (Supplementary Table 1). We note that the CAF and EC lines were the same throughout, and are not from the same patients as the cancer cells. Strikingly, we observed that migratory cell states emerge over time in tumoroids from certain patients and are not prevalent in others. (Fig. 4a). The migratory cells were observed from two patients that subsequently presented with recurrence, and had a shorter survival term than the other three patients (Supplementary Table 1). We generated single-cell transcriptome data from tumoroids from each patient (see Methods), analyzed cancer heterogeneity separately, integrated cancer cell data from each individual, visualized cells in a UMAP embedding, and identified markers for each cluster (Fig. 4b–d, Extended Data Fig. 7a-d, Supplementary Table 2). In the integrated analysis, we observed a diversity of cancer cell states and each individual contributed cells to all cell clusters (Extended Data Fig. 7e). Immunohistochemistry for Macrophage Migration Inhibitory Factor (MIF, enriched in cluster 0) showed a spatially distinct expression pattern in cancer cells at the periphery of the organoid (Fig. 4e). We compared each tumoroid cluster to bulk transcriptome datasets from different PDAC types (Classical A, B; Basal A, B) [30], and found that cell clusters within tumoroids could be classified based on signatures from primary cancer (Fig. 4f). Interestingly, we found that each tumoroid had cell states that expressed signatures of each of the different PDAC cancer types, and the proportions of these populations differ among patient tumoroids (Extended Data Fig. 7f). These data suggest that PDAC types classified from bulk measurements represent proportion differences among cancers, that the underlying cell states develop dynamically as a response to CAF stimuli, and that tumoroids might be used to recapitulate PDAC type proportions. Finally, we identify a multigene tumoroid migratory signature (TMS) that develops over time in tumoroids (Fig. 4g; Supplementary Table 2), induced in cancer cells by CAF, that positively correlates with shorter survival times (Fig. 4h,i; Extended Data Fig. 7g-i). We note that MIF and CEA-CAM6 are the two strongest indicators of the tumoroid migratory phenotype, and CEACAM6 has strong potential as a therapeutic target. High CEACAM6 expression is also associated with low cytotoxic T-cell infiltration [31]. Altogether, this modular developmental system, together with our analyses, provide a new inroad into the discovery of pancreatic cancer therapies.

**Fig. 4.**
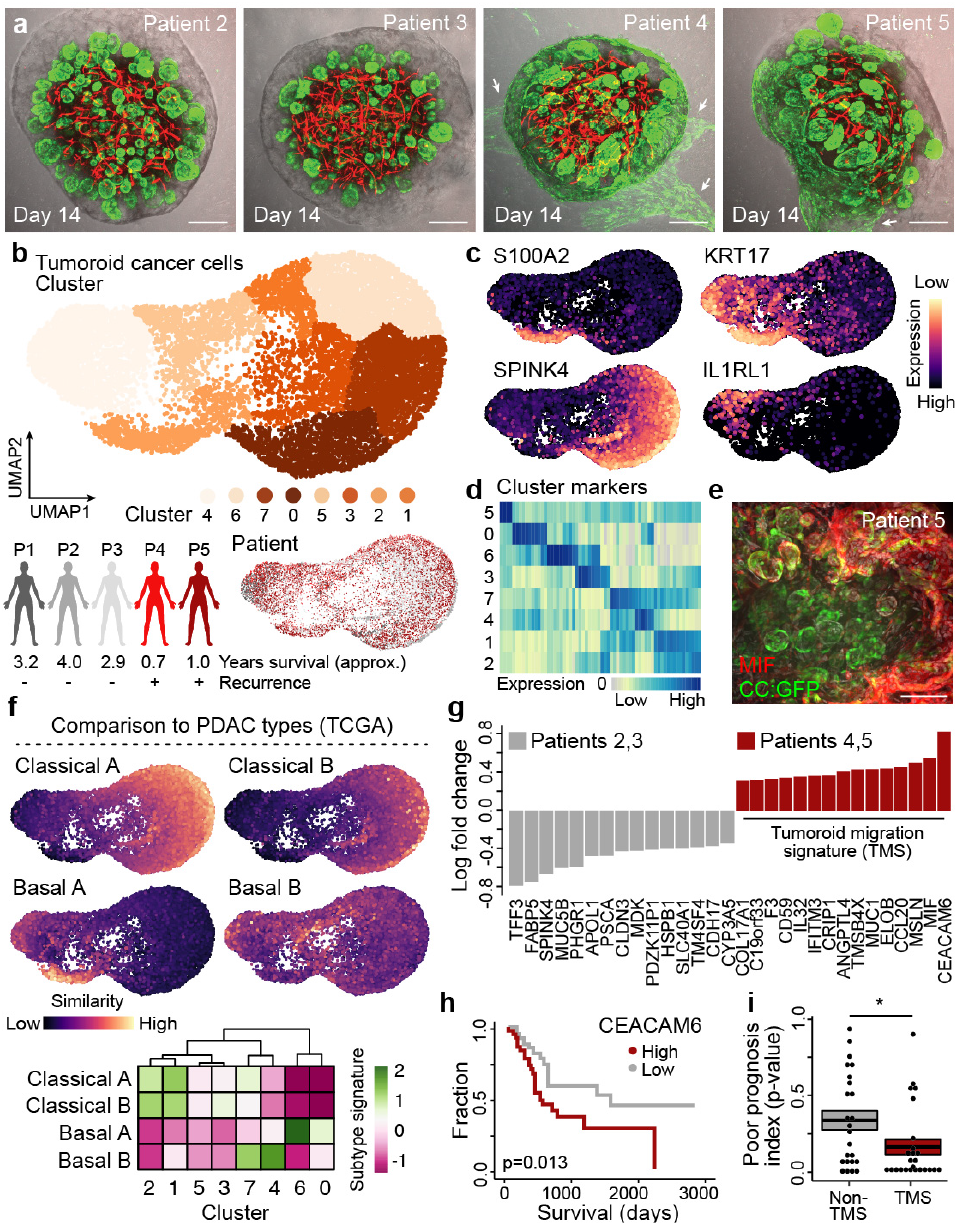
CAF-induced tumoroid migratory state signatures correlate with cancer prognosis. a) CCO cultures were established from additional PDAC patients and co-cultured with CAFs and ECs to generate tumoroids. Images show cancer cells and ECs stably transformed with EGFP and TdT expression cassettes, respectively. Arrows highlight the presence of migratory-like cells in certain patient tumoroids. Scale bar: 250um. b) Cancer cell heterogeneity analysis from scRNA-seq data from tumoroids from 5 patients. UMAP embedding is colored by cluster (top) and patient (bottom). Approximate overall survival time (in years) and recurrence status are noted for each patient. c,d) Feature plot (c) and heatmap (d) showing expression of cluster marker genes. e) Immunofluorescence staining for Macrophage Migration Inhibitory Factor (MIF) in organoids with cancer cells expressing GFP. Scale bar:100ul. f) Feature plots (top) and heatmap (bottom) showing scaled expression scores of PDAC subtype signatures in each tumoroid cancer cell or cluster, respectively. Data suggests co-existence of cancer sub-populations within the same tumoroid. g) Barplot shows log transformed fold changes of differentially expressed between cancer cells within tumoroids with and without migratory cell phenotypes. h) Kaplan–Meier (KM) plot showing that high expression of CEACAM6 in PDAC cancers from the cancer genome atlas (TGCA) dataset is associated with lower survival. i) Boxplot showing the p-value distribution of KM survival curve differences between high and low expression of genes in the tumoroid migratory signature (TMS) and non-migratory feature set.

We provide strong evidence that a stroma-rich in vitro PDAC microenvironment can induce cancer cells states that are relevant to primary PDAC tumor physiology. Most importantly, this system is dynamic and diverse phenotypic behaviors occur over time in response to intercellular communications. We highlight behaviors, interactions, and associated molecular states that can be manipulated for therapy development. In particular, CAFs activate during tumoroid formation and secrete matrix proteins, cytokines, and other cues that temporally impact cancer cell phenotypes, including metabolic adaptation and migratory escape from persistent cell stress. Strikingly, we observe migratory cell states in tumoroids from patients with worse clinical outcomes, and these tumoroids harbor potentially predictive signatures of poor prognosis. We stress that our experiments are from five patients, and future work is needed to explore the link between the patient and the tumoroid avatar. In addition, future work that incorporates macrophages, monocytes and other immune cell types will be required to fully recapitulate the dynamic interlineage signaling axes prevalent in PDAC tumor microenvironments. However, this tumoroid system is powerful in it’s modularity and reproducibility, providing an exciting new inroad into the mechanisms underlying fibroblast influence on cancer cell states.

## METHODS

### Establishment of cystic organoid and fibroblast cultures

The clinical specimens used to establish organoids and stromal cells were obtained from patients at the Kanagawa Cancer Center with informed consent after approval by the ethical review. Tumor tissue and healthy tissue were collected by surgical resection. The cancer cyst organoid (CCO) culture method from PDAC tumor specimens is briefly described below [14]. The surgical tissue is washed several times with Dulbecco’s phosphate buffered saline (DPBS). The tissue was finely c hopped u sing s urgical s cissors and a scalpel. The tissue was transferred to a 50 ml tube and washed again with DPBS. The washed tissue was digested with LiberaseTM (Roche) at 37°C for 40-60 minutes. Tissues were enzymatically treated and then washed with DMEM containing 10% fetal bovine serum (FBS, Sigma) to stop the enzymatic reaction. The obtained pancreatic cancer cells were embedded in growth factor reduced (GFR) Matrigel (Corning) and cultured in the following complete medium. DMEM/F12(Thermo), Primocine (1mg/ml, InvivoGen), GlutaMAX (1x, Invitrogen), 1x B27(1x, Invitrogen), Gastrin, N-acetyl-L-cysteine (1mM, Sigma), Nicotinamide (10mM, Sigma), A83-01(Tocris, 0.5uM), Noggin (Peprotech, 0.1ug/ml), R-Spondin1 (Peprotech, 100ng/ml), Wnt3A(RD, 50ng/ml), EGF (Peprotech, 50ng/ml), FGF10 (Peprotech, 100ng/ml). Y-27632 (Sigma, 10uM) was added for only one day after starting the organoid culture, and on the following day, the cells were cultured in a complete medium without Y-27632. The medium was changed 2 to 3 times a week. For establishment of fibroblasts, healthy pancreatic tissue and cancer tissue were treated with Liberase and the collected cells were washed with DPBS several times. Subsequently, cells were suspended in Mesenchymal stem cell growth media (MSCGM, Lonza) and seeded on a culture plate. The media was changed 2-3 times a week. All cells were cultured under 5% CO2 in 20% O2 at 37°C.

### Tumoroid culture method

To establish a stroma-rich pancreatic tumoroid, pancreatic CCO cells, fibroblasts and human umbilical vein endothelial cells (HUVECs) were separately expanded and cultured. Pancreatic CCO cells were incubated with Triple EX (Gibco) for 7 minutes, and fibroblasts and HUVECs were incubated for 3 minutes at 37°C to generate a cellular suspension. To stop the enzymatic reaction by Triple EX, the cells were washed with DMEM/F12 medium containing 10% FBS and 1% Penicillin-Streptomycin (P/S, Gibco). The obtained cells were counted separately, and then 3×104 cancer cells, 1.2×104 HUVECs, 8×104 fibroblasts were transferred to a tube coated with bovine serum albumin (1% BSA), mixed and centrifuged at 300 g. After removal of the supernatant, cell pellets were gently resuspended and 1.2 ×105 cells were then seeded in 96 well plates coated with 50% Matrigel (Corning). The three types of cells made cell-cell interactions with each other and showed self organisation during the period of 24-48 hours. The reconstituted stromal-rich pancreatic tumoroid were cultured with 50% Endothelial Cell Growth Media (Lonza) and 50% DMEM/F12 medium. Culture mediums were exchanged every 24 hours. Tumoroid were cultured under 5% CO2 in 20% O2 at 37°C.

### Generation of reporter lines

For live imaging, HUVECs were infected with retroviruses expressing Kusabira-Orange (KO) and cancer cells were infected with a lentivirus expressing enhanced green fluorescent protein (EGFP)[34]. Briefly, Human Embryonic Kidney (HEK) 283T cells were transfected with the retroviral vector pGCDNsam IRES-EGFP or KOFP (M. Onodera) for packaging at 293gag/pol (gp) and 293gpg (gp and VSV-G) to induce viral particle production. The culture supernatant of the retrovirus-producing cells was passed through a 0.45 mm filter (Whatman, GE Healthcare) and immediately used for infection. The firefly luciferase gene was subcloned into the CSII-EF-MCS-EGFP vector (RIKEN BRC) to generate the CSII-EF-Luc-IRES-EGFP construct. CSII-EF-Luc-IRES-EGFP plasmid and helper plasmid (293T cells were transfected with calcium phosphate using pCAG-HIVgp and pCMV-VSV-G-RSV-Rev, RIKEN BRC) to produce VSV-G pseudotyped lentivirus. The virus supernatant was recovered 46 hours after transfection, and filtered with a 0.45 µm filter. The virus supernatant was concentrated by ultracentrifugation.

### Whole-mount clearing and imaging

Tumoroids were washed several times with PBS and fixed with 200ul of 4% (wt/vol) paraformaldehyde (PFA). Tumoroids were incubated on a horizontal shaker at 4ºC for 24 hours. PFA was then completely removed and fixed tumoroids were washed several times with PBT buffer (0.1% Tween (vol/vol)). Tumoroid washing buffer (TWB: 100ml of PBS with 0.2 g of BSA and 0.1% Triton X-100) was added to the wells and incubated on a horizontal shaker at 4ºC for 1 day to block tumoroid. The next day, the blocking reagent was completely removed from the well, and then 100 ul of TWB with primary antibodies(1/100) was added to the wells and incubated on a horizontal shaker at 4ºC for 2 days. After immuno-labeling the tumoroid with the primary antibody, these reagents were removed from the wells, and then fresh TWB was added to the wells and was incubated on a horizontal shaker at 4ºC for 2 hours. This process was performed 3 times to completely remove the antibodies from organoids. After the tumoroid were sufficiently washed with TWB, 100ul of TWB with secondary antibodies(1/200) was added to the wells and incubated on a horizontal shaker at 4ºC for 1 day. After immuno-labeling the tumoroid with the secondary antibody, the secondary antibodies were removed from the wells and then fresh TWB was added to the wells and washed three times. Subsequently, 50ul of the fructose–glycerol clearing solution35 was added to the well and incubated on a horizontal shaker at 4ºC, overnight. Cleared organoids were placed on a glass slide or in a glass-bottom plate and imaged on a spinning disc confocal microscope (Olympus SpinSR10 spinning disk confocal super resolution microscope, objective x10,x20,x30,x40,x60). We used the following antibodies: anti-GFP (1:400; Abcam, ab13970), anti-Cytokeratin 7 (1:00; Agilent Technologies, M701829-2), anti-Cytokeratin 19 (1:100; Abcam, ab7754), anti-PPARG (1:100, Thermo Scientific, P A3-821A), a nti-Collagen I (1:100; Abcam, ab34710), anti-Fibronectin (1:100, Abcam, ab2413), anti-AKAP12 (1:100, Thermo Scientific, PA5-52281), anti-Trefoil Factor 3 (1:100, Abcam, ab108599), anti-Lipocalin-2 (1:100, Abcam, ab23477), anti-E-cadherin (1:100, RD Systems, AF748), anti-Syndecan-1 (1:100, Abcam, ab128936), anti-MIF (1:100, Abcam, ab187064), anti-Collagen III (1:100, Abcam, ab6310), anti-AGR2 (1:100, Sigma-Aldrich, HPA007912), anti-CEACAM6 (1:100, Thermo Scientific, MA5-29144), anti-MUC1 (1:100, Thermo Scientific, M A5-11202), D onkey a nti-Goat IgG (H+L) Highly Cross-Adsorbed Secondary Antibody, Alexa Fluor Plus 405 (1:200, Thermo Scientific, A48259), Donkey anti-Mouse IgG (H+L) Highly Cross-Adsorbed Secondary Antibody, Alexa Fluor Plus 555 (1:200, Thermo Scientific, A32773), Donkey anti-Rabbit IgG (H+L) Highly Cross-Adsorbed Secondary Antibody, Alexa Fluor Plus 647 (1:200, Thermo Scientific, A 32795), G oat A nti-Armenian hamster IgG HL (1:200,Abcam, ab173004), Goat anti-Chicken IgY (H+L) Cross-Adsorbed Secondary Antibody, Alexa Fluor Plus 488 (1:800, Thermo Scientific, A 32931), Molecular Probes DAPI (4’,6 Diamidino 2 Phenylindole, Dihydrochloride) (1:500, Thermo Scientific, D1306).

### SDC1 inhibition assays

For antibody treatment with anti-SDC1 (Abcam) on longterm cultured tumoroid, the co-cultured culture medium was completely removed from the wells and changed to 200ul of the co-cultured medium with 20ul anti-SDC1 added in the wells. The tumoroid were cultured under 5% CO2 in 20%O2 at 37°C. The antibody mixed medium was changed daily with a fresh one and the tumoroid had imaging after 5 days. A 200ul medium containing 20ul of isotype control antibody (Mouse IgG1, kappa monoclonal, Abcam) was used as a control medium.

### Single-cell RNA-seq experiments

All samples were dissociated to single cells by specific enzymatic treatment. The cultured medium for stroma-rich tumoroid was removed from the wells and tumoroids washed three times with 1xDPBS. The tumoroids were collected in 5 ml tubes, after the DPBS was completely removed from the tube, TrypLE™ Select (Thermo) was added and incubated at 37ºC for 8 minutes [36]. After the incubation step, tumoroids were further dissociated by trituration. This incubation and trituration process was repeated 3 times to obtain a single cell suspension. The enzymatic dissociation was stopped by addition of cold BE-PBS (Cold PBS 1 ml with 0.04% BSA / (0.1 mM EDTA)) and remaining cellular clumps were removed by using 70um and 40um strainers. Fibroblasts and HUVECs were cultured on a 10 cm dish and dissociated to a single cell suspension using the same procedure as described above. Single cell suspensions were adjusted to an appropriate concentration to obtain approximately 2000-10000 cells per lane of a 10x Genomics microfluidic Chip G. Libraries were generated using 10x Genomics 3’ Gene Expression Kit (v3.1), following recommended protocol, and subsequently sequenced on NextSeq500, using 28-9-0-91 Read Configuration, as recommended by for Single Index libraries.

### Single-cell RNA-seq data preprocessing

CellRanger (v3.1.0, 10x Genomics) was used to extract unique molecular identifiers, cell barcodes, and genomic reads from the sequencing results of 10x Chromium experiments. Then, count matrices, including both protein coding and non-coding transcripts, were constructed aligning against the annotated human reference genome (GRCh38, v3.0.0, 10x Genomics). In order to remove potentially damaged or unhealthy cells and improve data quality, the following filtering steps were performed in addition to the built-in CellRanger filtering pipeline. Cells associated with over 20,000 transcripts, usually less than 1% of the total number of samples, were removed. Cells associated with a low number of transcripts (<1% of the total number of samples) were removed. Cells with over 15% of mitochondrial transcripts were removed. Transcripts mapping to ribosomal protein coding genes were ignored. Cells with <800 unique transcripts (<1% of the total number of samples) were removed together with transcripts detected in less than 10 samples.

### Normalization with Seurat

For normalization and variance stabilization of each scRNAseq experiment’s molecular count data, we employed the modeling framework of SCTransform in Seurat v3 [37]. This procedure overcomes the need for heuristic steps by performing a more effective normalization, strongly removing technical effects from the data while preserving biologically relevant sources of heterogeneity. In brief, a model of technical noise in scRNA-seq data is computed using ‘regularized negative binomial regression’. The residuals for this model are normalized values that indicate divergence from the expected number of observed UMIs for a gene in a cell given the gene’s average expression in the population and cellular sequencing depth. The residuals for the top 2,000 variable genes were used directly as input to computing the top 100 Principal Components (PCs) by PCA dimensionality reduction through the RunPCA() function in Seurat. Corrected UMI, which are converted from Pearson residuals and represent expected counts if all cells were sequenced at the same depth, were log-transformed and used for visualization and differential expression (DE) analysis.

### Doublet removal with DoubletFinder

For each scRNA-seq experiment DoubletFinder [38] was used to predict doublets in the sequencing data. Its workflow c an b e b roken u p i nto 4 m ajor s teps. F irst, generate artificial doublets from existing scRNA-seq data by merging randomly selected cells. Second, pre-process merged real-artificial d ata. T hird, p erform P CA a nd u se t he PC distance matrix to find e ach c ell’s p roportion o f a rtificial k nearest neighbors (pANN). For this step we restricted the dimension space to the top 30 PCs. Fourth, rank order and threshold pANN values according to the expected number of doublets. Optimal input parameters for doublet estimation and removal were selected through ROC analysis across pN-pK parameter sweeps for each scRNA-seq dataset; pN describes the proportion of generated artificial doublets while pK defines t he P C n eighborhood s ize. I n o rder to achieve maximal doublet prediction accuracy, mean-variance normalized bimodality coefficient ( BCmvn) w as leveraged. This provides a ground-truth-agnostic metric that coincides with pK values that maximize AUC in the data. DoubletFinder is mostly sensitive to heterotypic doublets but not to homotypic ones. To overcome this we consider doublet number estimates based on Poisson statistics with homotypic doublet proportion adjustment assuming 1/50,000 doublet formation rate the 10x Chromium droplet microfluidic cell loading.

### Data integration with Cluster Similarity Spectrum (CSS)

Individual datasets, after preprocessing and doublet removal, were aggregated according to specific criterias ( e.g. patient id, timepoint, culture condition) and went through an additional step of mild processing in order to mitigate technical confounding factors, which also served as means for selection of a set of meaningful 2,000 most variable global genes prior to data integration. Integration of different conditions (cell lines and timepoints) was performed using the log-normalized corrected UMI count data. We used the first 3 0 P Cs a nd t he P earson r esiduals t o i ntegrate the different timepoints (or cell lines) in the datasets using the Cluster Similarity Spectrum method (CSS) [39]. In brief, clustering is applied to cells within each sample label separately and similarities, by Spearman correlation, of one cell to those clusters are calculated and normalized. To obtain a two-dimensional (2D) representation of the data we performed Uniform Manifold Approximation and Projection (UMAP [39,40] using RunUMAP() with default parameters on the CSS matrix. Integrated datasets were then clustered according to the shared neighborhood graph on lower dimensional space using the Louvain algorithm41 through either the Seurat function FindClusters() with resolution 0.2.

### Pseudotime reconstruction

PCA is an eigenvector-based multivariate analysis that defines a new orthogonal coordinate system that optimally describes variance in a dataset. It learns a linear transformation where the PCs form an orthogonal basis for the features that are uncorrelated [42]. By construction, this transformation can encode the original data in a latent (lower dimensional) space concentrating much of the signal into the first few principal components and achieve a higher signal-to-noise ratio while minimizing the total squared reconstruction error. Given its strength, we thus sought at using PCA to learn time-dependent variability in our tumoroid system and optimally describe heterogeneity in the scRNA-seq time course data by reconstructing a differentiation trajectory for each cell type in the 2D PCA space. Cells were subsequently aligned along that trajectory. This was done for the time-course data of the multi-lineage tumoroid cells derived from patient 1 in co-culture with CAF and endothelial cells; cells in other culture conditions or derived from different patients were then projected to that space.

### CAF-tumor communication

To investigate ligand-receptor (LR) mediated cell-cell communications during cancer progression in our multi-lineage tumoroids, we focused on the signals exchanged between CAF and cancer cells. For this analysis we extracted genes labeled as either ligands or receptors from curated databases43 and performed differential expression (DE) tests between CAF and cancer cells in order to gain insights into specific molecules involved and retrieve directional information about the signal exchange. Among all significant LR pairs, we focused first on CAF to cancer signaling, thus considering CAF as the signal source, expressing ligands, and cancer as the signal target, expressing receptors. We then looked also at signals being delivered by cancer to CAF receptors. In order to model dynamic communication during cancer progression we emphasized LR pairs characterized by significant incremental expression change along pseudotime alignment. This analysis identified ligand-receptor (LR) pairs which significantly co-expressed along CAF-cancer trajectories, and therefore potentially mediated the communications between cell populations. To prioritize most important signaling molecules we first modelled the gene expression as smooth functions over pseudotime. In particular, we used a univariate penalized cubic regression spline basis smooth [44] defined by a set of 80 knots spread evenly through the covariate (pseudotime) values. They were penalized by the integrated square second derivative cubic spline penalty, modified to shrink towards zero at high enough smoothing parameters. Next, we used cosine similarity to summarize the interaction (binding) strength for every LR pair; higher the cosine similarity is the stronger the signaling through the given LR pair. Uncertainties (standard errors) resulting from spline fits at each knot were propagated through the computation of the cosine similarity to report a confidence estimate on its measure.

### Functional enrichment analysis

To understand mechanisms underlying phenotypes in our data, differentially expressed genes were analyzed for cancer hallmark enrichment using one-sided hypergeometric testing. P-values were adjusted for multiple testing hypotheses by the Bonferroni method and only enrichment results below a 5% significance level threshold were considered. The hallmarks are a collection of curated gene sets, within MSigDB, refined to convey a specific b iological s tate o r p rocess a nd display coherent expression [45]. Cell populations were evaluated for over representation or change in biologically related functional gene sets. For this analysis we only considered hallmarks consisting of sets with more then 10 but less then 300 mapped genes.

### Gene regulatory network inference in tumoroids

To gain biological insights into mechanisms driving time-dependent cellular heterogeneity in pancreatic cancer, we resorted to utilizing the Single-Cell rEgulatory Network Inference and Clustering (SCENIC) workflow [ 16]. This is a computational method to infer gene regulatory networks (GRNs) and cell types from single-cell RNA-seq data. In brief, we initialized SCENIC options by selecting the latest versions (v9) of two motif annotation datasets, 500 base pairs (bp) upstream and 100 bp downstream of transcription start site (TSS) as well as 10k bp centered TSS, on human genome (hg38) and 20 processing units; otherwise default parameters. To infer potential transcription factor targets, we imputed Pearson residuals of the most variable genes expressed in at least 10% of the cells and built a co-expression network via GENIE346 with default parameters. GENIE3 is a Random Forest based method capable of detecting nonlinear dependencies. In order to distinguish activation from repression we took advantage of the Spearman correlation between transcription factors and respective target genes. Finally, the activity of the inferred GRNs was computed by aggregating the expression of the target genes within each single cell. We then identified t op variable G NRs by assessing their activity along the pseudotime.

### Bulk RNA-seq data processing

Pancreas adenocarcinoma (PAAD) related samples from The Cancer Genome Atlas (TCGA) were downloaded via the Genomic Data Common (GDC) website. The R package TGAbiolinks was used to connect to GDC data transfer tool client and GDC API in order to query and download the raw gene expression profiles, metadata and available clinical data for 177 tumor samples as well as 4 normal pancreatic tissue samples. The data was filtered b y r emoving subsequent occurrences of probes matching the same gene symbol as well as probes matching no known genes at all. Raw HTSEQ counts data was then normalized for sequencing depth using estimateSizeFactors() and variance-stabilized through regularized-logarithm transformation to remove spurious effects from aberrant gene counts with rlog() function in DESeq2 [47]. For the regularized-logarithm transformation the blind parameter was set to false. This, to ensure that variables in the design formula will not contribute to the expected variance-mean trend of the experiment; otherwise default parameters were used.

### Survival analysis

Individual samples were divided into higher and lower categories based on normalized expression of the gene of interest. A quartile cutoff based on expression was considered to group the samples into two categories. The long-term survival probability was analyzed by utilizing the Kaplan-Meier survival plot. Log-rank test [48], as implemented within the survdiff() function, was applied to assess the difference between the survival cohorts. Analysis of survival time was performed using the survival package in R statistical software.

### Differential expression analysis

Gene differential expression (DE) between distinct cell populations in scRNA-seq data was assessed by performing Wilcoxon rank sum tests and auROC analysis as implemented in Presto [49] package in R. Log-transformed corrected UMIs were used as input for the DE statistical tests, and genes were called differentially expressed if associated adjusted p-value (Bonferroni method) was lower than 0.05, AUC value was above 0.6 and log fold change was greater than 0.15. In addition we also set thresholds on detection rates of DE genes. In particular, a given gene was assigned as over-expressed in the analyzed group if it was detected in at least 30% of the samples of that group while the detection rate in the background samples was at most 70% of the detection rate of the analyzed group. To probe association between genes and invasive phenotype we performed within-cluster DE tests between different cell lines. The resulting specificity score is then an aggregate count of the times a given gene was found to be a top 5 up-regulated genes for a specific phenotype. Differential expression analysis on bulk RNA-seq data was performed in accordance with the DESeq2 pipeline on row counts with DESeq() function and default parameters. Genes were then considered up-regulated if they were assigned an adjusted p-value below 0.05 and a log fold change greater than 0.5.

## AUTHOR CONTRIBUTIONS

RO developed the protocol with support from YU, HT. RO generated tumoroids and performed IHC experiments. RO, BG performed single-cell RNA-seq experiments and analyzed data with support from QY, DP, ZH, AM, MSa, MSe, RO. Patient material was provided by SM, YM, TY. RO, HT, BT and JGC designed the study and RO, BG, DP, AM, and JGC wrote and edited the manuscript.

## ACKNOWLEDGEMENTS

We thank the Camp and Treutlein labs, and Keisuke Sekine for helpful discussions. This work was supported in part by Chan Zuckerberg Initiative DAF, an advised fund of the Silicon Valley Community Foundation CZF2019-002440 (J.G.C., B.T.), the Swiss National Science Foundation (Project Grant-310030_84795, J.G.C.; Project Grant-310030_192604, B.T.), and the National Center of Competence in Research Molecular Systems Engineering (B.T.). R.O. was supported by the Japanese Society for the Promotion of Science (JSPS).

## DATA AVAILABILITY

Raw sequencing data will be deposited into ArrayExpress. Processed data and the VCF files for demultiplexing will be deposited in Mendeley Data.

## Supplementary Figures

**Supplementary Fig. 1.**
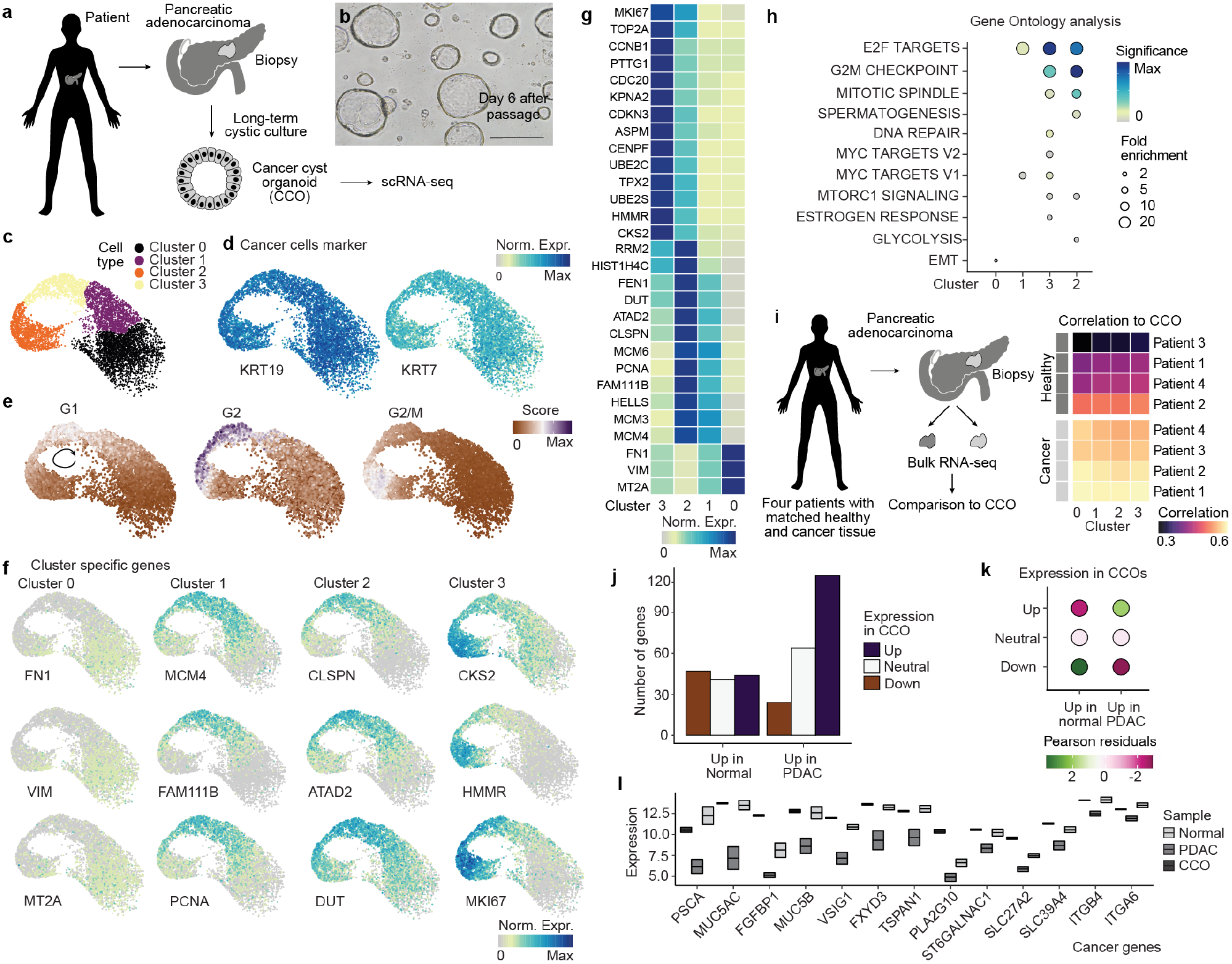
Single-cell transcriptome analysis of pancreatic cancer cyst organoid cultures and comparison to primary cancer tissue. a) Cancer cyst organoid (CCO) lines were established from PDAC patient biopsies. CCOs were propagated over multiple passages, and after day 7 of culture post-passage, CCOs were subjected to scRNA-seq. Scale bar:400um. b) Brightfield image of CCOs in a 3D matrigel culture 6 days after passage. c) UMAP cell embedding of CCO scRNA-seq data colored by cluster. d-f) Feature plots of PDAC cancer cell markers Keratin (KRT19 and KRT17), correlation to cell cycle states (e) and cluster markers (f). g) Heatmap showing normalized cluster marker expression. h) Gene ontology analysis of genes enriched in each CCO cluster. Circles are colored based on significance (High p-value in blue, low in yellow/gray) and sized by fold enrichment. i) CCO pseudo-bulk samples, obtained through aggregation of single cells according to their respective cluster memberships, were compared to bulk transcriptome data from healthy and cancer tissue from the TCGA cohort. Heatmap shows correlation (high, bright yellow; low, dark colors) of each cluster to the different patient specimens. j) Barplots show the number of genes that are differentially expressed between normal and PDAC tissue and are up-regulated (up), downregulated (down), or not differentially expressed (neutral) in CCOs compared to primary pancreas cells. k) Dotplot shows similarity between CCO and pancreatic cancer signatures when compared to healthy pancreatic tissue (chi-squared p-value 1.6e-07). l) Boxplots show the expression distribution of pancreatic cancer markers as comparison between TCGA healthy and cancer samples and our CCO culture.

**Supplementary Fig. 2.**
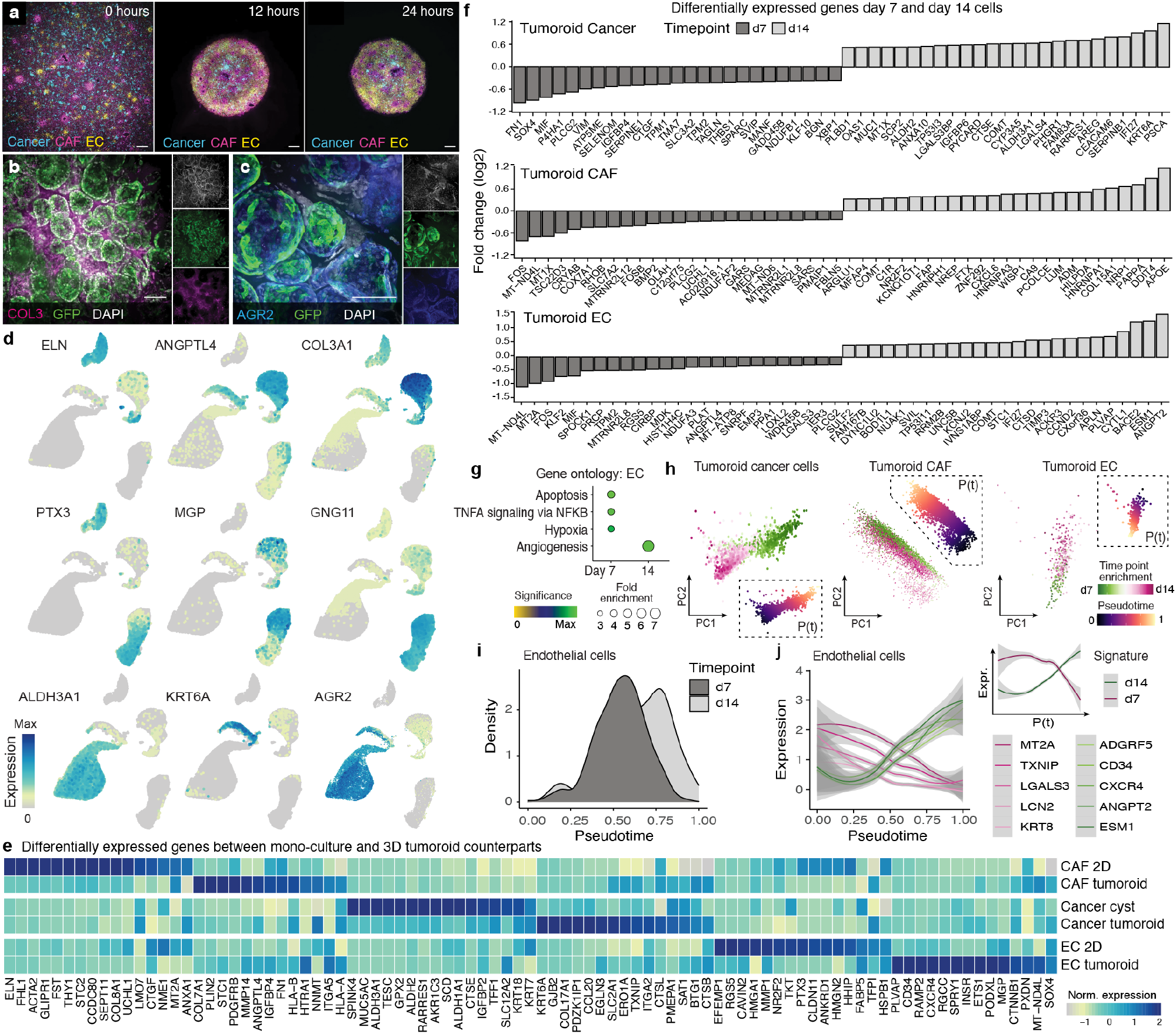
Comparison between 2D monoculture and 3D tumoroid cells and tumoroid time points. a) Cancer cells (teal blue), cancer associated fibroblasts (CAFs, pink), and endothelial cells (ECs, yellow) expressing a reporter 0, 12, and 24 hours after co-culture. Scale bar:10ul. b-c) Whole-mount immunohistochemistry on cleared tumoroids probing Collagen 3 (COL3, b) and Anterior gradient 2 (AGR2, c). Cancer cells stably expressing EGFP; nuclei are marked with DAPI. Scale bar:100ul. d) Feature plots showing expression of representative cluster and cell type marker genes from single-cell transcriptome data generated from tumoroids and input cells. e) Heatmap showing genes that are differentially expressed between tumoroid cell types and their input counterparts. f) Barplots show transformed fold change (log 2) between day 7 (dark grey) and day 14 (light grey) tumoroid cancer (top), CAF (middle), and EC (bottom) cells. g) Gene ontology analysis of genes enriched in day 7 and day 14 tumoroid endothelial cells. h) Principal component (PC) analysis was used to establish pseudotemporal ordering of cancer, CAF, and endothelial cells, and plots show cells colored by pseudotime. i) Density plots showing proportion of tumoroid endothelial cells along the inferred pseudotime. j) Pseudotemporal expression profile of genes differentially expressed (DE) between tumoroid endothelial cells at day 7 and day 14. Inset shows cumulative expression profiles of the DE genes.

**Supplementary Fig. 3.**
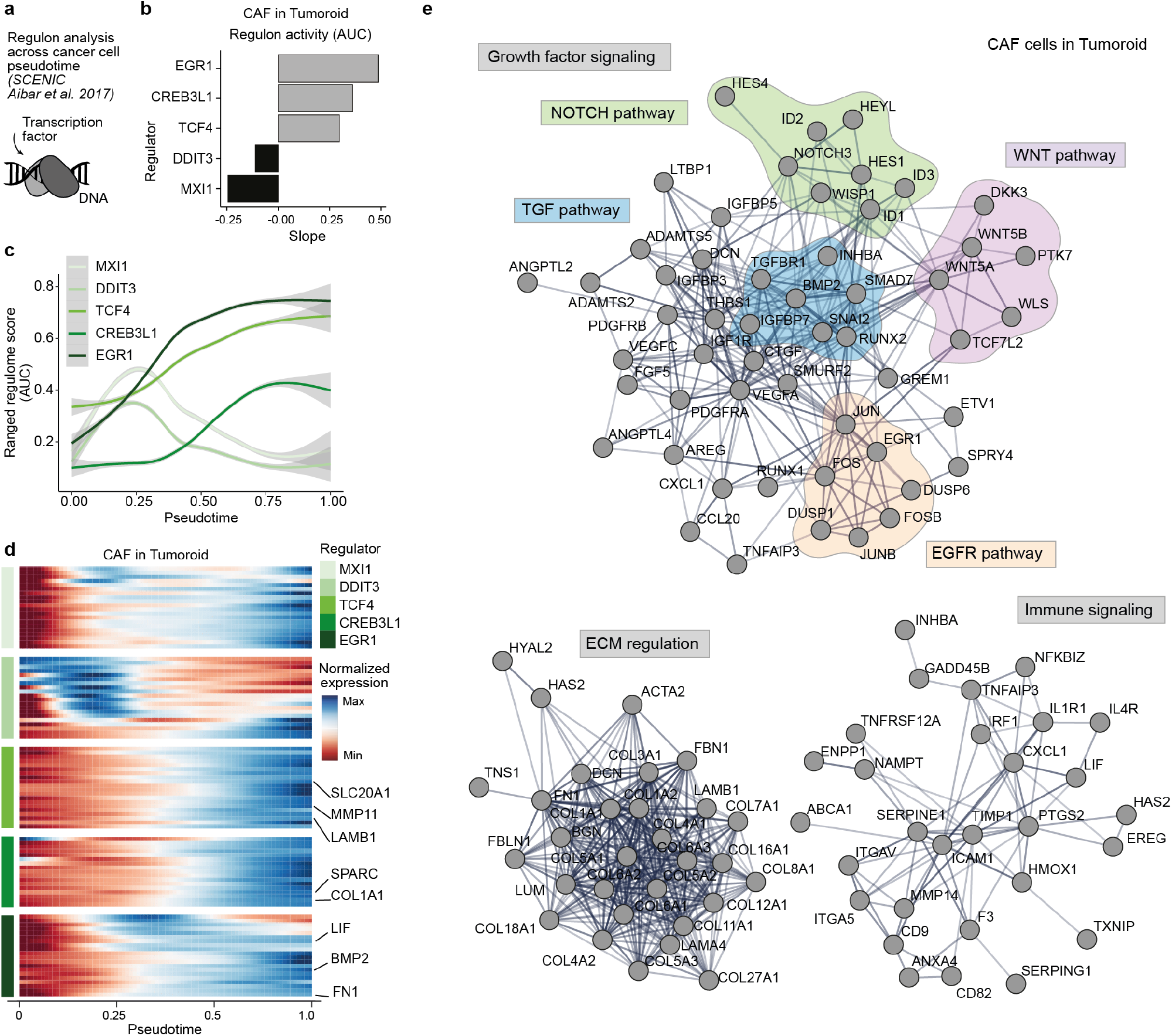
Regulon analysis of Tumoroid CAFs. a) SCENIC was used to infer regulatory network scores for tumoroid CAFs. b) The expression slope over tumoroid CAF pseudotime is plotted for the top regulators based on area under the curve (AUC) metrics from SCENIC. c) Line plots show the ranged regulon score for each of the top regulators over tumoroid CAF pseudotime. d) Heatmap shows expression of predicted targets of each major regulator over tumoroid CAF pseudotime. e) Protein-protein interactome from STRING database with pathway annotation for up-regulated networks for CAF pseudotemporally regulated genes.

**Supplementary Fig. 4.**
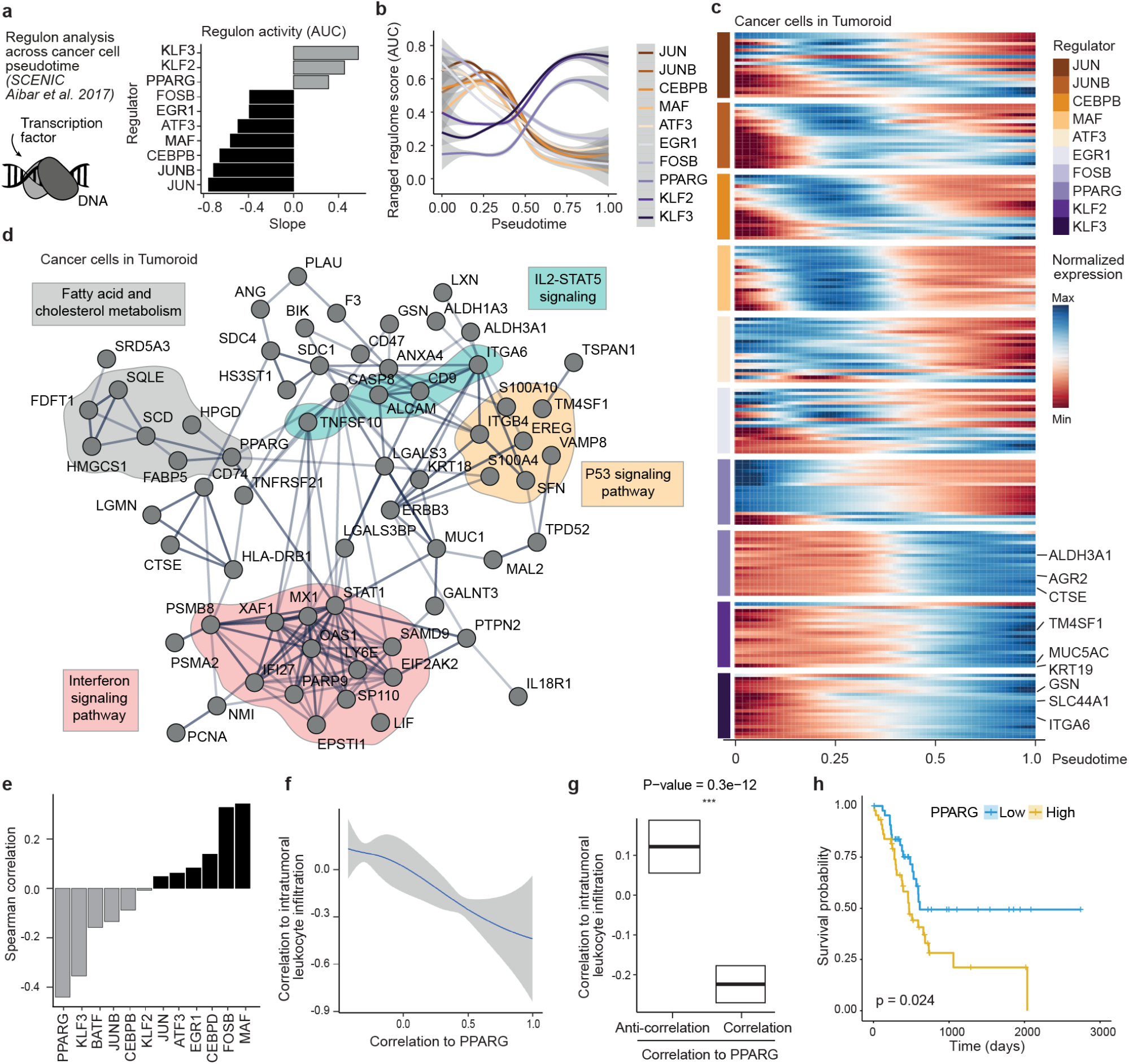
Regulon analysis of Tumoroid cancer cells. a) SCENIC workflow was used to infer regulatory network scores for tumoroid cancer cells (left). The expression slope over tumoroid pseudotime is plotted for the top regulators based on area under the curve (AUC) metrics from SCENIC (right). b) Line plots show normalized pseudotemporal expression of central regulators. c) Heatmap shows expression of predicted targets of each major regulator over tumoroid cancer cell pseudotime. d) Proteinprotein interactome from STRING database with pathway annotation for up-regulated networks. e) Spearman correlation score between transcription factors (TFs) of dynamic regulons and percentage of tumor infiltrating immune cells in TCGA pancreatic cancer sample cohort. TFs show a wide range of values, and note that PPARG registers the highest negative correlation score. f) Correlation plot of PPARG expression with the percentage of infiltrating intratumoral immune cells g) Differential association of positively versus negatively PPARG correlated genes with percentage of intratumoral immune cell infiltration is statistically significant. h) Kaplan-Meier curve showing significant association of high PPARG expression with poor prognosis in the TCGA PDAC cohort.

**Supplementary Fig. 5.**
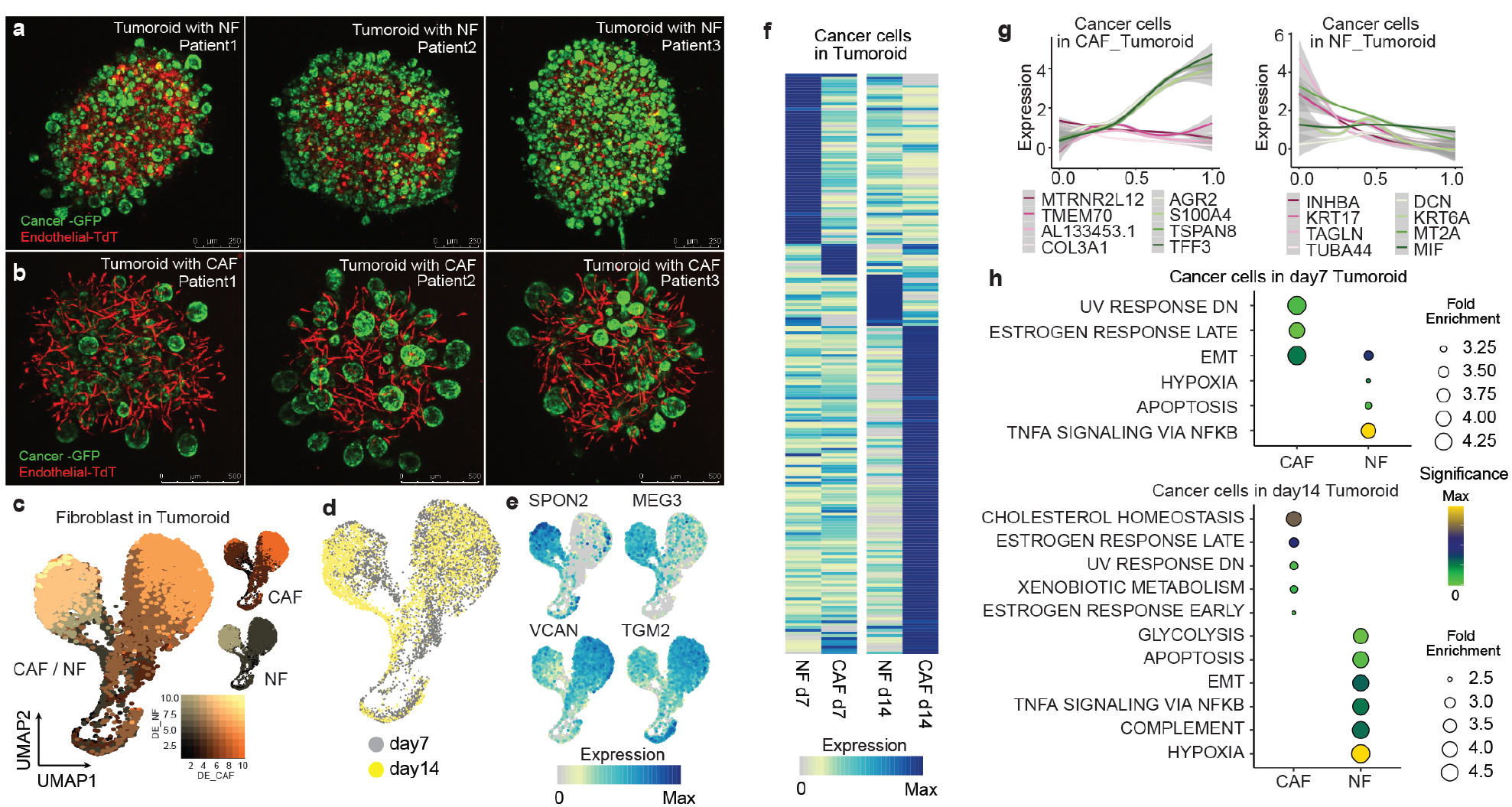
Comparison of NF and CAF tumoroids. a-b) Images show tumoroids generated with normal fibroblasts (NFs, a - upper row) or cancer associated fibroblasts (CAFs, b - lower row) with endothelial cells labeled with TdTomato and cancer cells labeled with EGFP. Scale bar:250um. c-e) UMAP cell embedding from Fig. 2 showing normal and cancer associated fibroblasts colored by respective transcriptional signatures (c), tumoroid culture timepoint (d), or expression feature (e). f) Heatmap shows expression profiles of differentially expressed (DE) genes, at each timepoint, between cancer cells in tumoroids with NFs or CAFs. g) Pseudotemporal expression pattern of representative genes in cancer cells from NF and CAF tumoroids. h) Hallmark enrichment analysis in cancer cells within the different tumoroid types at 7 (top) and 14 (bottom) days of co-culture. Data shows how cancer cells within the CAF tumoroid change state and acquire new metabolic footprints, while cancer cells in contact with normal fibroblasts remain largely stable over the entire co-culture period.

**Supplementary Fig. 6.**
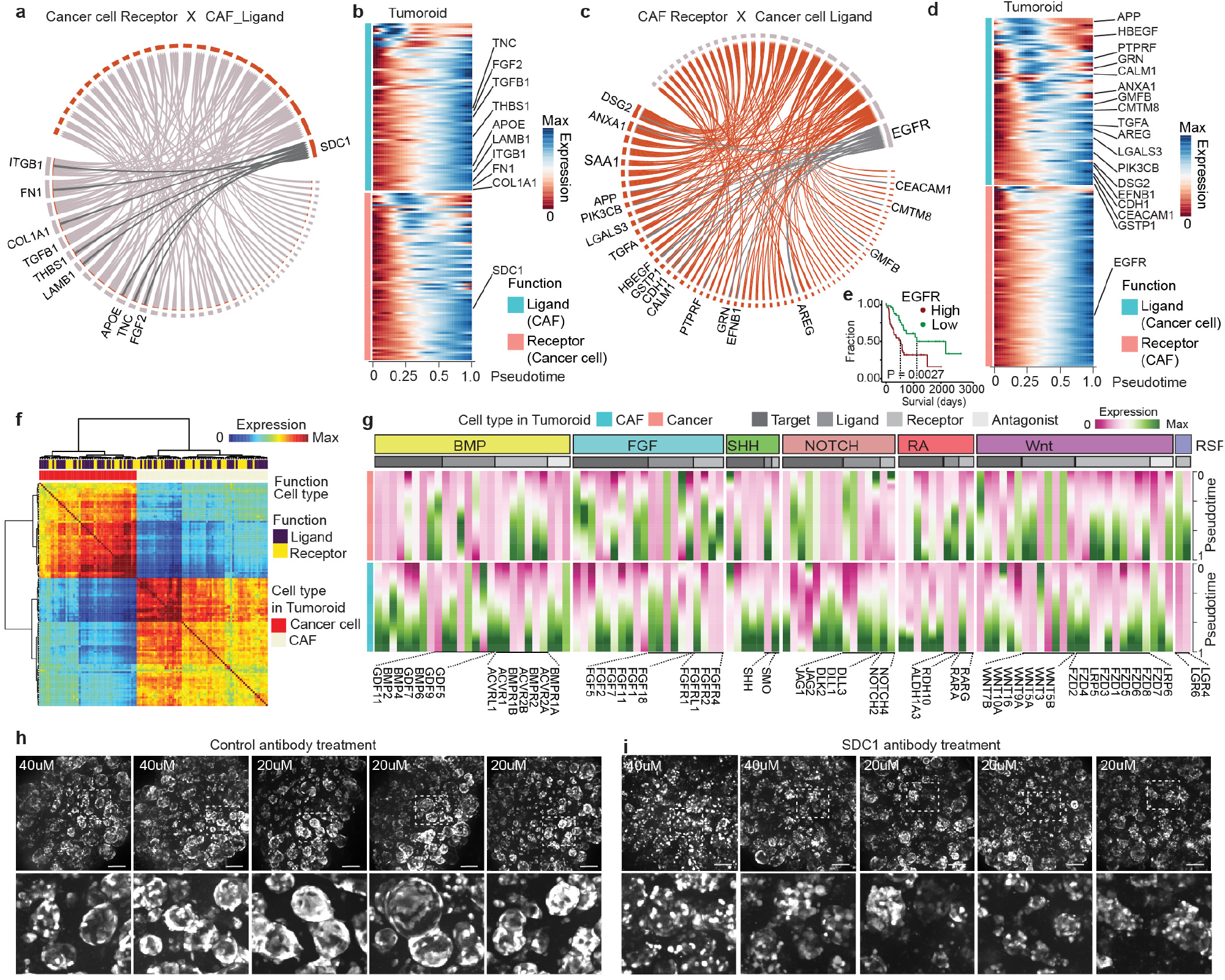
Expression of receptors and ligands in tumoroid CAF and cancer cells. a) Ribbon plot representing communication between cancer associated fibroblasts (source) and cancer cells (target) highlighting SDC1 interactions. b) Expression of CAF-specific ligands and cancer-specific receptors along respective pseudotemporal trajectories. c) Ribbon plot representing communication between cancer cells (source) and CAFs (target) highlighting EGFR interactions. EGFR is inferred to be a major hub for collecting cancer stimuli and up-regulated during fibroblast activation. d) Expression of cancer-specific ligands and CAF-specific receptors along respective pseudotemporal trajectories. e) Survival analysis on TCGA pancreatic cancer cohort links EGFR overexpression to worse outcome. f) Cross-correlation of ligands’ and receptors’ expression patterns reveals distinct repertoires associated with each tumoroid cell type. g) Signaling molecules from curated annotations33 of major developmental pathways are dynamically expressed along CAF and cancer trajectories within tumoroids. h) GFP reporter expression in cancer cells on day 14 tumoroids incubated with isotype control (left) or SDC1 (right) antibodies. Scale bar:100um

**Supplementary Fig. 7.**
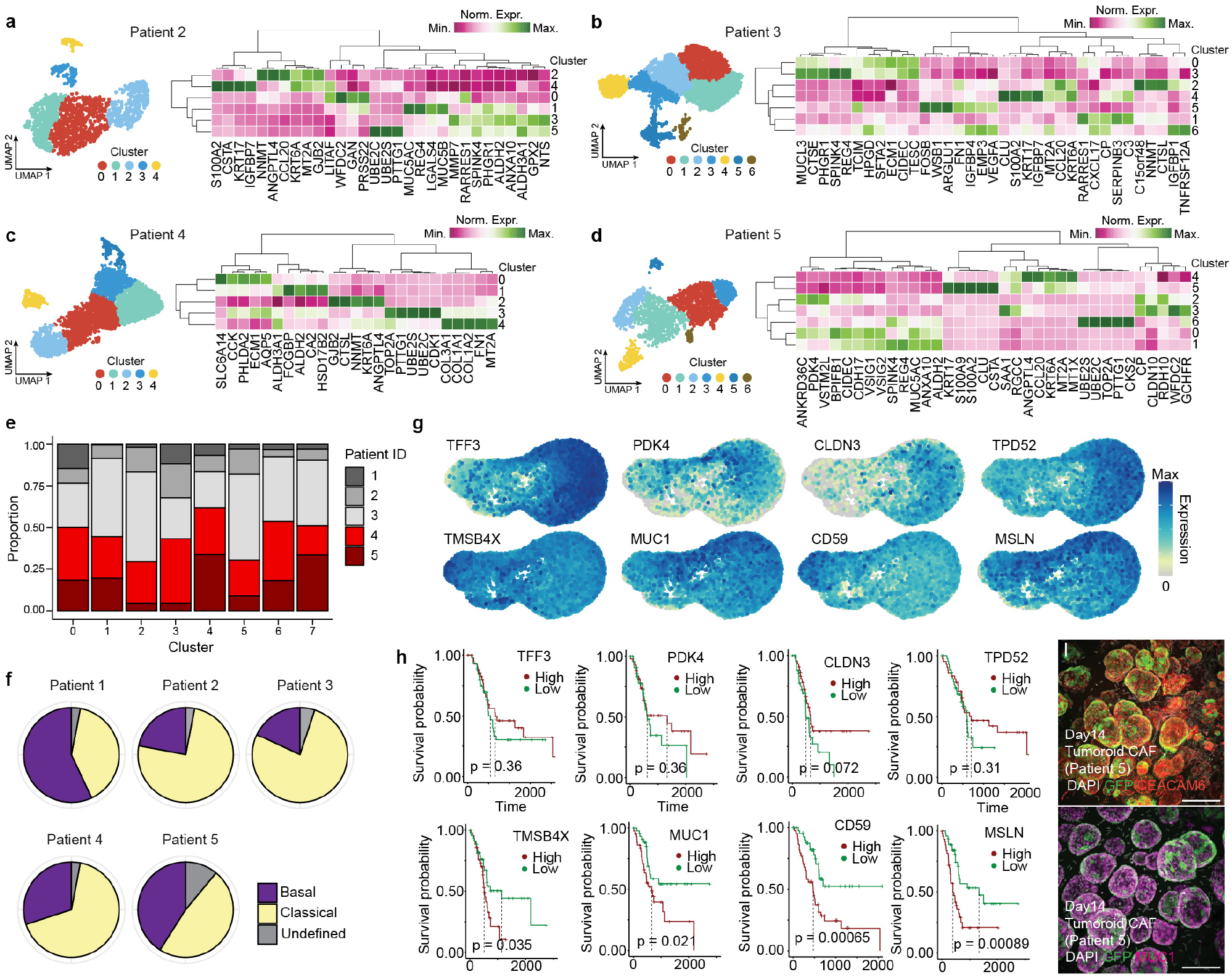
Heterogeneity analysis of tumoroid cancer cells from six patients. a-d) Heterogeneity analysis in tumoroid cancer cells for each patient derived line showing selection of marker genes for cultures with no migratory cells (a,b) and cultures with migratory cells (c,d). Insets display individual UMAP embeddings color-coded by cluster. Note that the cluster colors are not comparable across samples. e) Proportion of cells in each cluster that are derived from each patient from the integrated heterogeneity analysis presented in Fig. 4. f) Pie chart shows the proportion of cells from each patient tumoroid classified into PDAC subtypes. The data shows how different subtypes co-exists within each patient-derived tumoroid. g) Feature plots on the integrated UMAP from Fig. 4 showing expression of genes enriched in tumoroids without (top row) and with (bottom row) migratory cells. h) Kaplan–Meier (KM) plot showing the relationship of gene expression in the Tumor Migration Signature (TMS, bottom) or in the non-TMS (top) and the survival times. Data is from the cancer genome atlas (TGCA). i) Immunofluorescence staining for CEACAM6 (red, top) and MUC1 (pink, bottom) on day 14 tumoroids from Patient 6. Cancer cells stably express GFP, DAPI marks nuclei (white). Scale bar:100um.

## Notes

### Competing Interest Statement

The authors have declared no competing interest.

